# Spatial patterning of P granules by RNA-induced phase separation of the intrinsically-disordered protein MEG-3

**DOI:** 10.1101/073908

**Authors:** Jarrett Smith, Deepika Calidas, Helen Schmidt, Tu Lu, Dominique Rasoloson, Geraldine Seydoux

## Abstract

RNA granules are non-membrane bound cellular compartments that contain RNA and RNA binding proteins. The molecular mechanisms that regulate the spatial distribution of RNA granules in cells are poorly understood. During polarization of the *C. elegans* zygote, germline RNA granules, called P granules, assemble preferentially in the posterior cytoplasm. We present evidence that P granule asymmetry depends on RNA-induced phase separation of the granule scaffold MEG-3. MEG-3 is an intrinsically disordered protein that binds and phase separates with RNA *in vitro*. *In vivo*, MEG-3 forms a posterior-rich concentration gradient that is anti-correlated with a gradient in the RNA-binding protein MEX-5. MEX-5 is necessary and sufficient to suppress MEG-3 granule formation *in vivo*, and suppresses RNA-induced MEG-3 phase separation *in vitro*. Our findings support a model whereby MEX-5 functions as an mRNA sink to locally suppress MEG-3 phase separation and drive P granule asymmetry.

**HIGHLIGHTS:**

- The intrinsically-disordered protein MEG-3 is essential for localized assembly of P granules in *C. elegans* zygotes.
- MEG-3 binds RNA and RNA stimulates MEG-3 phase separation.
- The RNA-binding protein MEX-5 inhibits MEG-3 granule assembly in the anterior cytoplasm by sequestering RNA.

## Introduction

RNA granules are concentrated assemblies of RNA and RNA-binding proteins that form without a limiting membrane in the cytoplasm or nucleoplasm of cells (Courchaine, Lu et al. 2016). RNA granules are ubiquitous cellular structures and several classes of cytoplasmic RNA granules have been described, including stress granules, P bodies, neuronal granules and germ granules (Anderson and Kedersha 2006). Cytoplasmic RNA granule components typically exchange rapidly between a highly concentrated pool in the granule and a more diffuse, less concentrated pool in the cytoplasm (Weber and Brangwynne 2012). In addition to RNA-binding domains, proteins in RNA granules often contain prion-like, low complexity, or intrinsically-disordered regions (IDRs) (Courchaine, Lu et al. 2016). In concentrated solutions, IDRs spontaneously demix from the aqueous solvent to form liquid droplets (liquid-liquid phase separation or LLPS) or hydrogels (Li, Banjade et al. 2012, Weber and Brangwynne 2012, Elbaum-Garfinkle, Kim et al. 2015, Guo and Shorter 2015, Lin, Protter et al. 2015, Nott, Petsalaki et al. 2015). Like RNA granules *in vivo*, proteins in LLPS droplets and hydrogels exchange with the solvent (Kato, Han et al. 2012, Li, Banjade et al. 2012, Elbaum-Garfinkle, Kim et al. 2015, Lin, Protter et al. 2015). These findings have suggested that LLPS or reversible gelation drives the assembly of RNA granules *in vivo* (Guo and Shorter 2015).

In cells, RNA granule assembly is regulated in space and time. For example, stress granules assemble within seconds of exposure to toxic stimulants that require the temporary removal of mRNAs from the translational pool (Anderson and Kedersha 2006). In eggs, germ granules assemble in the germ plasm, a specialized area of the cytoplasm that is partitioned to the nascent germline during the first embryonic cleavages (Voronina, Seydoux et al. 2011). How phase separation, a spontaneous process *in vitro*, is regulated *in vivo* to ensure that RNA granules form at the correct place and time is not well understood.

The germ (P) granules of *C. elegans* are an excellent model to study the mechanisms that regulate granule assembly (Updike and Strome 2010). For most of development, P granules are stable perinuclear structures, but in the transition from oocyte-to-embryo, P granules detach from the nucleus and become highly dynamic (Pitt, Schisa et al. 2000, Wang, Smith et al. 2014). As the oocyte is ovulated in the spermatheca, P granules disassemble and release their components in the cytoplasm. After fertilization, P granule proteins reassemble into dynamic granules that undergo repeated cycles of assembly and disassembly in synchrony with cell division. Live imaging in the 1-cell zygote has revealed that these cycles are spatially patterned along the anterior-posterior axis of the embryo: granule assembly is favored in the posterior and granule disassembly is favored in the anterior (Brangwynne, Eckmann et al. 2009, Gallo, Wang et al. 2010). By the first mitosis, P granules are found exclusively in the posterior cytoplasm together with other germ plasm components. As a result, P granules are segregated to the posterior germline blastomere P_1_ and excluded from the anterior somatic blastomere AB.

P granule asymmetry is under the control of the PAR polarity network which divides the zygote into distinct anterior and posterior domains (Motegi and Seydoux 2013). The PAR-1 kinase is enriched in the posterior cytoplasm and restricts the RNA-binding protein MEX-5 (and its redundant homolog MEX-6) to the anterior cytoplasm (Griffin, Odde et al. 2011). MEX-5 and MEX-6 in turn restrict P granules to the posterior (Schubert, Lin et al. 2000, Gallo, Wang et al. 2010). In *mex-5 mex-6* double mutants, P granule still undergo cycles of assembly and disassembly but these are no longer patterned along the anterior-posterior axis, and small granules remain throughout the cytoplasm (Gallo, Wang et al. 2010). An attractive hypothesis is that MEX-5 blocks phase separation of P granule components in the anterior cytoplasm (Brangwynne, Eckmann et al. 2009, Lee, Brangwynne et al. 2013). The mechanism of MEX-5 action and the critical P granule component(s) regulated by MEX-5, however, are not known.

P granule assembly in zygotes requires several P granule proteins, including the RNA-binding protein PGL-1 (and its redundant paralog PGL-3) and the intrinsically-disordered protein MEG-3 (and its redundant paralog MEG-4) (Hanazawa, Yonetani et al. 2011, Wang, Smith et al. 2014). PGL-1 and PGL-3 are RGG domain proteins that self-associate and recruit other RNA-binding proteins to the granules, including the GLH family of RNA helicases (Updike and Strome 2010, Hanazawa, Yonetani et al. 2011). MEG-3 and MEG-4 are redundant, serine-rich proteins that bind to PGL-1 *in vitro* and are essential for P granule assembly in embryos. In zygotes lacking *meg-3* and *meg-4*, PGL-1 and GLH-2 form transient assemblies that do not segregate asymmetrically and are not maintained in later stages (Wang, Smith et al. 2014). In this study, we use a combination of *in vivo* and *in vitro* experiments to examine the contribution of the MEX, MEG and PGL proteins to P granule asymmetry. We show that MEG-3/4, but not PGL-1/3, are essential for granule asymmetry, and that MEX-5 localize MEG-3 in a posterior-rich gradient. We demonstrate that MEG-3 is an RNA-binding protein that is stimulated by RNA to undergo phase separation. MEX-5 is sufficient to block MEG-3 phase separation *in vitro* and to block MEG-3 granule formation *in vivo*. Our findings are consistent with a model whereby MEX-5 sequesters RNA to inhibit MEG-3 condensation in the anterior cytoplasm.

## RESULTS

### Hierarchical regulation of P granule assembly

To determine the genetic hierarchy that controls granule asymmetry, we first compared the distributions of MEX-5, MEG-3, PGL-1 and GLH-1 during the earliest stages of zygote polarization. We monitored MEX-5, MEG-3 and GLH-1 localization using tagged alleles generated by genome editing (Methods, Table S1) and PGL-1 localization using a polyclonal antibody that recognizes PGL-1 (Strome and Wood 1983).

Before polarization, MEX-5, MEG-3, PGL-1 and GLH-1 were all distributed evenly throughout the cytoplasm. MEX-5 appeared mostly diffuse in the cytoplasm, whereas MEG-3, PGL-1, and GLH-1 appeared both diffuse and enriched in many small (<1 micron diameter) granules (Fig 1A). During symmetry breaking (pronuclear formation and migration), MEX-5 and MEG-3 began to redistribute into opposing cytoplasmic gradients along the long axis of the zygote (anterior-posterior axis) with MEG-3 beginning to form large (∼1 micron) granules in the posterior (Fig 1A). Total levels of MEG-3 do not change during this period, consistent with redistribution of existing MEG-3 protein from anterior to posterior (Fig 1B, Fig S1A). In contrast to MEX-5 and MEG-3, PGL-1 and GLH-1 remained uniformly distributed during symmetry breaking. By mitosis, all proteins were localized, with MEX-5 in the anterior cytoplasm and MEG-3, PGL-1 and GLH-1 in large granules in the posterior cytoplasm (Fig 1A).

**Figure 1.**
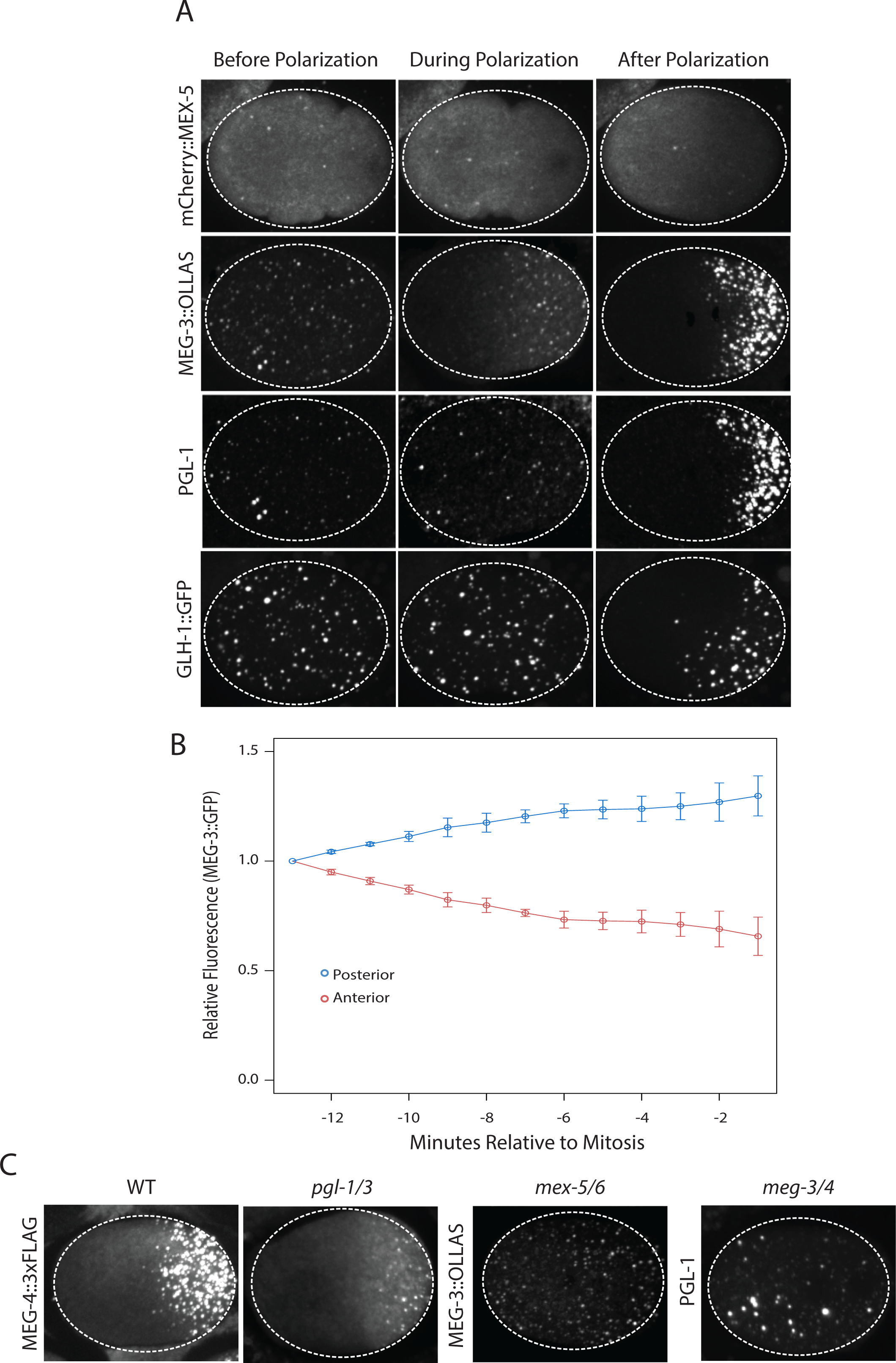
Localization of P granule proteins during zygote polarization. A. Photomicrographs of live wild-type (mCherry::MEX-5 and GLH-1::eGFP) or fixed *meg-4* zygotes (MEG::3 OLLAS and PGL-1) at three different stages: before polarization (pronuclear formation), during polarization (pronuclear migration) and after polarization (mitosis). *meg-4(ax3052)* zygotes were coimmunostained for MEG-3::OLLAS (anti-OLLAS, Novus Biological), and PGL-1 (K76, DSHB). In this and subsequent figures, dashed circles represent outline of embryo, embryos are oriented with anterior to the left and posterior to the right, and all embryos are ∼50 µM long. *meg-*4 is required redundantly with *meg-3* for P granule assembly, and each is sufficient to support localized granule assembly (Wang, Smith et al. 2014). B. MEG-3::meGFP levels in anterior and posterior halves of the zygote during polarization. Values represent average fluorescence intensity over time (relative to initial levels) in the anterior (red) and posterior (blue). Averages come from values measured from 3 embryos. Error bars represent standard deviation of the mean. C. Photomicrographs of fixed zygotes immunostained for FLAG, OLLAS or PGL-1. *pgl-1/3* zygotes were derived from *pgl-3(bn104)* hermaphrodites treated with *pgl-1* RNAi. *mex-5/6* zygotes were derived from wild-type hermaphrodites treated with *mex-5* and *mex-6* RNAi. *meg-3/4* zygotes were derived from *meg-3(ax3055); meg-4(ax3052)* hermaphrodites.

To determine the interdependence of these localizations, we examined the effect of removing MEX-5/6, MEG-3/4 or PGL-1/3 using RNA-mediated interference (RNAi) or genetic mutants. In zygotes derived from mothers treated with double-stranded RNA against *mex-5* and *mex-6* (*mex-5/6(RNAi)* zygotes), the MEG-3 gradient did not form and MEG-3 and PGL-1 granules remained uniformly distributed throughout the cytoplasm (Fig 1C, (Gallo, Wang et al. 2010). In *meg-3meg-4* double mutant embryos, the MEX-5 gradient was unaffected but PGL-1 granules did not segregate (Wang, Smith et al. 2014, Fig. 1C). In *pgl-1/3(RNAi)* zygotes, the MEG-3 and MEG-4 gradients were unaffected and MEG-3/MEG-4 granules formed in the posterior as in wild-type, except that the granules appeared smaller (Fig 1C, Fig S1B, (Wang, Smith et al. 2014). These analyses suggest that MEX-5 and MEX-6 regulate granule asymmetry by localizing MEG-3 and MEG-4 to the posterior, which in turn are required to localize PGL-1 and PGL-3. PGL-1 and PGL-3 are not required to localize MEG-3 or MEG-4, but contribute to the increase in size of MEG-3/4 granules at mitosis.

### MEX-5 is necessary and sufficient to suppress MEG-3 granule formation

Using *mex-5* transgenes, Griffin *et al.* 2011 showed that formation of the MEX-5 gradient requires phosphorylation of serine 404 in the C-terminus of MEX-5 by the kinase PAR-1. To determine whether the MEX-5 gradient is required to pattern MEG-3, we mutated serine 404 to alanine (S404A) at the *mex-5* locus using CRISPR/Cas9-mediated genome editing (Paix, Folkmann et al. 2015). We introduced the S404A mutation in two strains: one where the *mex-5* locus had been previously tagged with mCherry to monitor MEX-5 localization, and one where MEG-3 had been previously tagged with GFP to monitor MEG-3 localization (Table S1). As expected, we found that mCherry::MEX-5(S404A) failed to form a gradient during zygote polarization and remained uniformly distributed (Fig 2A). Using the MEG-3::GFP strain, we found that zygotes derived from mothers homozygous for *mex-5(s404a)* (*mex-5(S404A)* zygotes), MEG-3::GFP did not form a gradient or granules. Instead, MEG-3::GFP remained uniformly distributed in the cytoplasm throughout the 1-cell stage (Fig 2B). We conclude that MEX-5 is sufficient to suppress the formation of MEG-3 granules throughout the cytoplasm.

**Figure 2.**
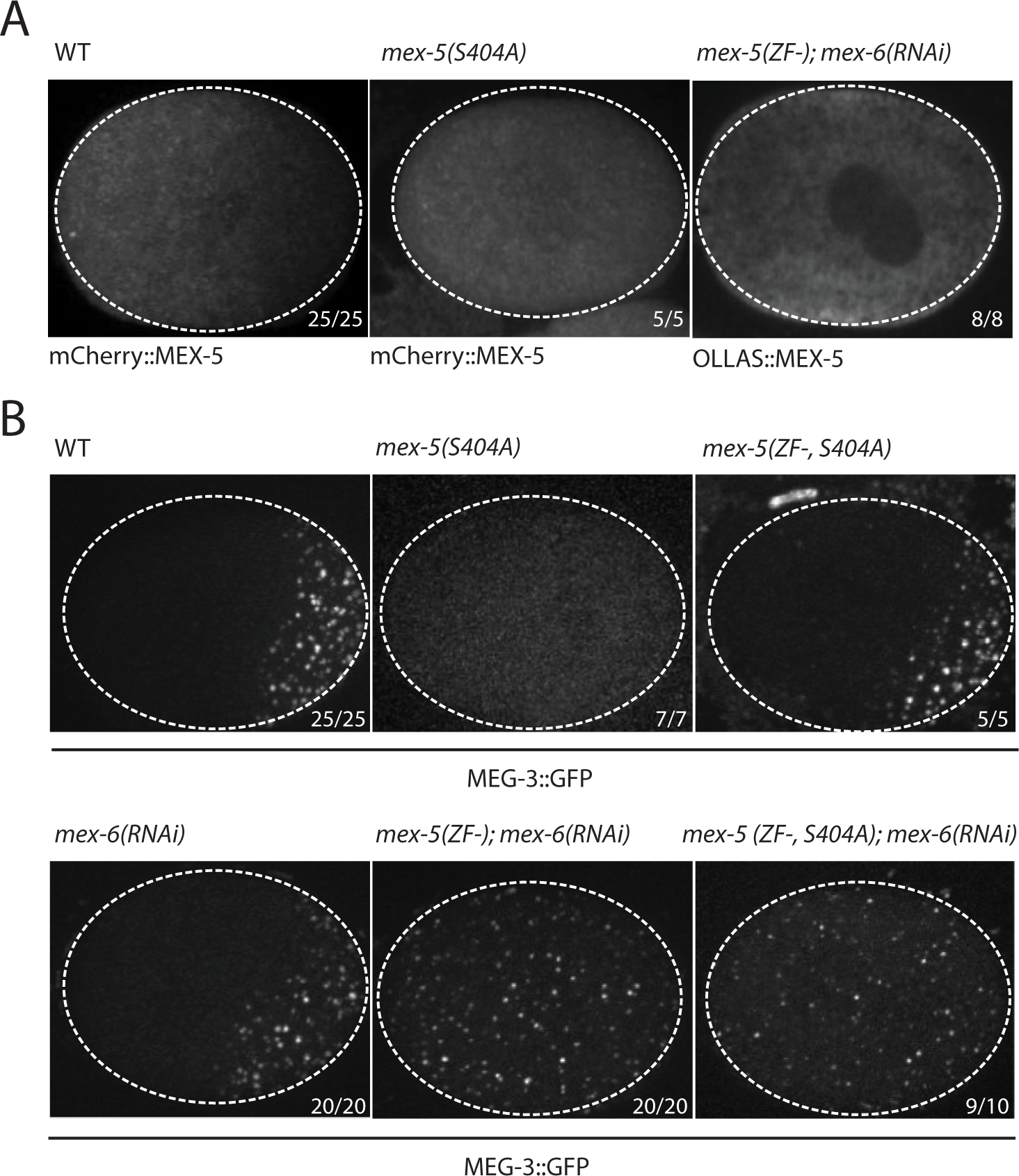
MEX-5 is necessary and sufficient to disassemble MEG-3 granules *in vivo*. A. Photomicrographs of live [wild-type and *mex-5(S404A)*] or fixed [*mex-5(ZF-);mex-6(RNAi)*] zygotes at pronuclear meeting to show MEX-5 localization. Wild-type MEX-5 is in an anterior-rich gradient, whereas MEX-5(S404A) and MEX-5(ZF-) are uniformly distributed. Numbers indicate number of zygotes exhibiting phenotype shown / total number of zygotes examined. B. Photomicrographs of live embryos expressing MEG-3::GFP. Genotypes at the *mex-5* locus are as indicated. Numbers are as in A. In 1/10 *mex-5(ZF-, S404A); mex-6(RNAi)* zygotes, MEG-3 granules were asymmetric possibly due to incomplete depletion of MEX-6.

### MEX-5 RNA binding is required to suppress MEG-3 granule assembly

The MEX-5 RNA-binding domain is comprised of two zinc fingers that bind with high affinity to poly-U stretches. Pagano *et al.* 2007 have shown mutation of a single amino acid in each finger (R247E and K318E) reduces MEX-5 binding affinity for poly-U by 35-fold. To determine whether suppression of granule assembly by MEX-5 requires high affinity RNA-binding, we introduced R247E and K318E (hereafter referred to as ZF-) at the *mex-5* locus by CRISPR/Cas9 genome editing in the mCherry::MEX-5 and MEG-3::GFP lines. Like MEX-5(S404A), mCherry::MEX-5(ZF-) did not form a gradient and was uniformly distributed in zygotes (Fig 2A). In contrast to *mex-5(S404A)* zygotes, however, *mex-5(ZF-)* zygotes assembled posterior MEG-3 granules as in wild-type (data not shown). RNAi depletion of *mex-6*, the non-essential *mex-5* paralog (Schubert, Lin et al. 2000), in this background yielded zygotes that assembled MEG-3 granules throughout the cytoplasm, as in *mex-5/6(RNAi)* zygotes (Fig 2B). These observations suggest that *mex-5(ZF-)* is a loss-of-function allele. The loss of *mex-5* activity was not due to reduced expression as MEX-5(ZF-) was expressed at the same level as wild-type MEX-5 (Fig S2).

To determine whether high-affinity RNA binding is also required for MEX-5(S404A) ability to suppress MEG-3::GFP granule assembly throughout the cytoplasm, we introduced the S404A mutation by genome editing into *mex-5(ZF-)* hermaphrodites. We found that *mex-5(ZF-,S404A)* zygotes assembled posterior MEG-3::GFP granules, as is observed in *mex-5(ZF-)* and wild-type zygotes. Depletion of *mex-6* by RNAi in this background yielded zygotes with uniform MEG-3::GFP granules, as expected for a *mex-5* loss-of-function phenotype (Fig 2B). We conclude that suppression of MEG-3 granule assembly by MEX-5 depends on MEX-5’s ability to bind RNA with high affinity.

### MEG-3 binds RNA *in vitro*

Unlike MEX-5, MEG-3 does not have a recognizable RNA-binding domain. MEG-3 contains a long predicted intrinsically-disordered region (IDR) at its N-terminus (aa1-550) followed region with lower predicted disorder (aa550-862) [IUPRED predictions, (Dosztanyi, Csizmok et al. 2005)]. To determine whether MEG-3 binds RNA, we expressed and purified as His-tagged fusions full length MEG-3, MEG-3(aa1-544) (hereafter referred to MEG-3_IDR_) and MEG-3(aa545-862) (hereafter referred to MEG-3_Cterm_) (Fig S3A). We tested each for binding to poly-U30 RNA using electrophoretic mobility shift assays (EMSA) and fluorescent polarization (FP) assays (Pagano, Farley et al. 2007). EMSA experiments revealed that MEG-3 and MEG-3_IDR_ interact with poly-U30 RNA to form complexes that migrate as a discrete band during electrophoresis (Fig 3A). Using FP, we calculated the apparent dissociation constant (K_d,app_) of MEG-3 for poly-U30 to be ∼32 nM, similar to that of MEX-5 (K_d,app_ = ∼29 nM) (Pagano, Farley et al. 2007). MEG-3_IDR_ also bound RNA but with ∼15-fold lower affinity (K_d,app_ = ∼460nM). MEG-3_Cterm_ did not bind RNA significantly by EMSA or in the FP assay (K_d,app_ > 3000 nM) (Fig 3, Fig S3B). We conclude that MEG-3 binds RNA with high affinity and that this activity resides primarily within the MEG-3 IDR, although high affinity binding also requires the MEG-3 C-terminus.

**Figure 3.**
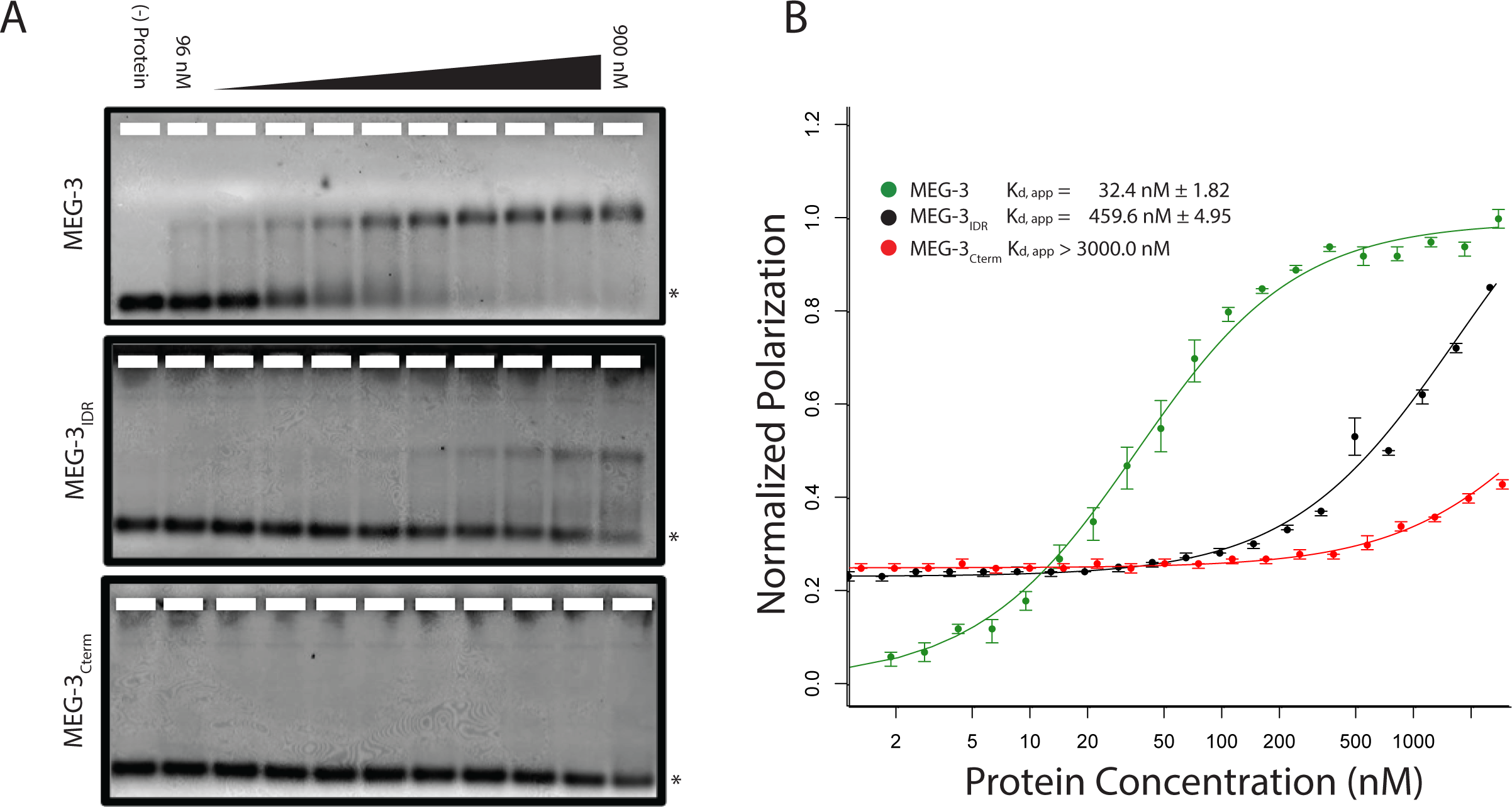
MEG-3 Binds RNA *in vitro*. A. Binding of MEG-3 to poly-uridine 30 (poly-U30) is shown by electrophoretic mobility shift assay (EMSA) using fluorescein-labeled poly-U. EMSAs are shown for (top to bottom) full length MEG-3, MEG-3_IDR_, and MEG-3_Cterm_. Unbound poly-U30 is denoted by an asterisk (*). B. Fluorescence Polarization of poly-U30 by MEG-3. Fluorescence polarization values normalized relative to saturation are shown for full length MEG-3 (green), MEG-3_IDR_ (black), and MEG-3_Cterm_ (red). A fit of the polarization as function of protein concentration is plotted and used to calculate the given K_d,app_. Error bars report S.E.M. Expanded graphs are shown in Fig S3B.

To begin to explore the specificity of MEG-3 RNA binding, we challenged MEG-3/poly-U30 complexes with increasing concentrations of competitor RNAs and examined their behavior by EMSA. We found that poly-U, and to a lesser extent poly-A, were effective competitors, but not poly-C or poly-G (Fig S3C). These observations suggest that MEG-3’s affinity for RNA is affected by nucleotide composition.

### MEG-3 and MEG-3_IDR_ phase separate *in vitro*

Concentrated (<1 µM) solutions of RNA-binding proteins containing IDRs spontaneously phase separate when switched from high to low salt (150 mM NaCl) (Elbaum-Garfinkle, Kim et al. 2015, Lin, Protter et al. 2015, Nott, Petsalaki et al. 2015). We were not able to maintain high concentrations of MEG-3 or MEG-3_IDR_ in solution in the presence of high salt (Methods). Therefore to examine the phase separation properties of MEG-3, we used concentrated (100-320 µM) preparations maintained in 6M urea, diluted these into aqueous buffer (150 mM NaCl) and used light microscopy to immediately observe the mixture (Method). We found that MEG-3 and MEG-3_IDR_ readily formed phase-separated condensates within 10 min at room temperature. Control proteins (BSA and MBP) subjected to the same treatment did not phase separate (data not shown). MEG-3 condensates were observed across a range of protein concentrations (0.5 µM to 5 µM) and became larger and more abundant with increasing protein concentration (Fig S4A). MEG-3 and MEG-3_IDR_ behaved similarly to each other, except that in the low concentration range (< 5 µM), MEG-3 formed more condensates than MEG-3_IDR_, and in the high concentration range (> 5 µM) MEG-3_IDR_ tended to form larger condensates (Fig S4A).

RNA can stimulate the phase transition of IDR proteins that bind RNA (Guo and Shorter 2015). To determine the effect of RNA on MEG-3 phase separation, we added poly-U30 RNA to the phase separation buffer before diluting in MEG-3. We found that 0.1 µM poly-U30 was sufficient to increase the number of visible condensates especially at low MEG-3 protein concentrations (<1 µM) (Fig 4A and Fig S4A). Higher concentrations of RNA increased the number of MEG-3 condensates even further. For a given concentration of poly-U30, MEG-3 formed more condensates than MEG-3_IDR_ (Fig 4A and Fig S4A). Addition of sub-stochiometric amounts of fluorescently tagged poly-U30 confirmed that the RNA phase separates with MEG-3 (Fig S4B). We conclude that MEG-3 and MEG-3_IDR_ have an intrinsic propensity to phase separate that can be stimulated by RNA.

**Figure 4.**
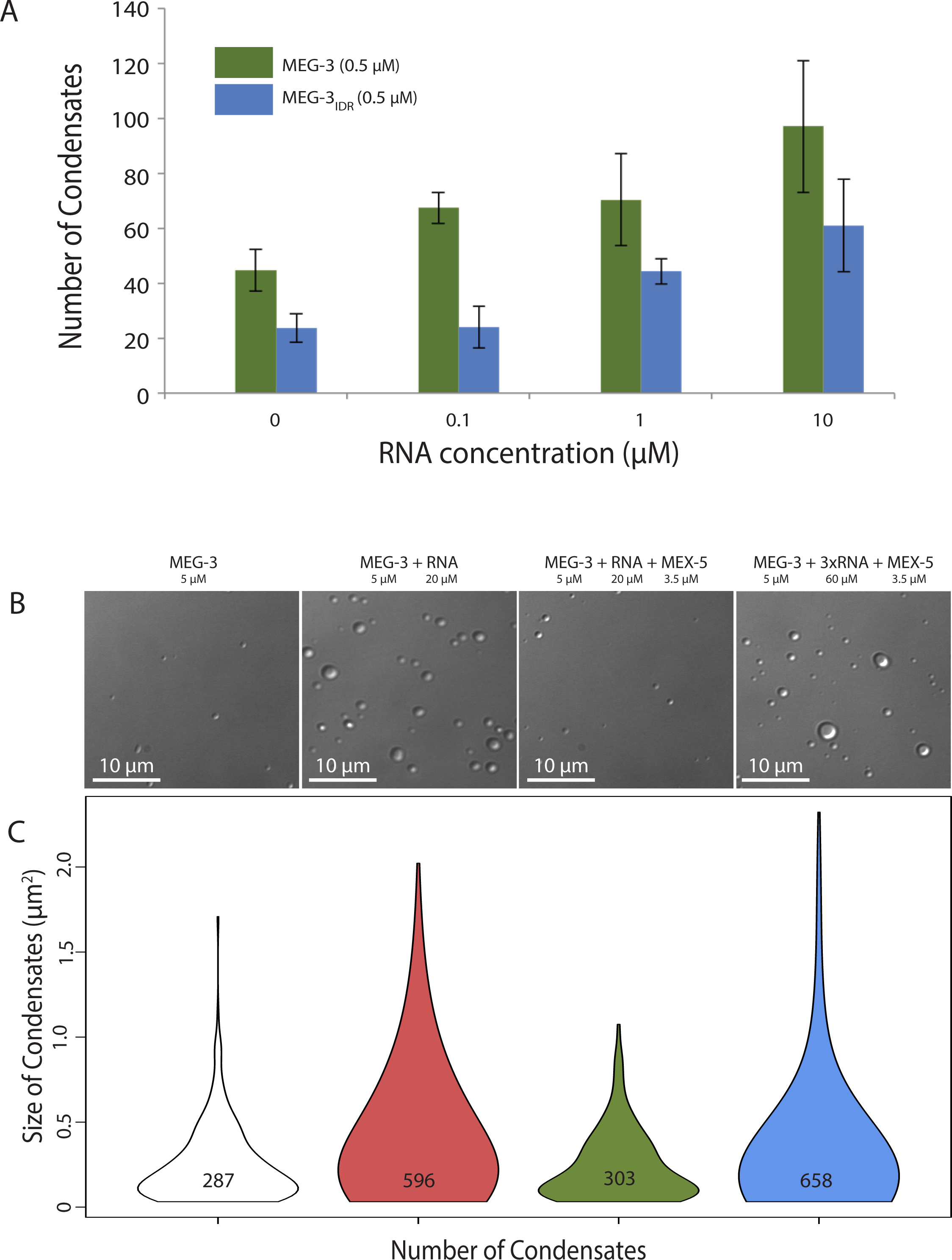
Stimulation of MEG-3 phase separation by RNA. A. Bar graph showing the number of condensates formed by 0.5 µM MEG-3 or MEG-3_IDR_ in the presence of increasing poly-U30. Error bars indicate S.E.M from three technical replicates for each condition. B. Photomicrographs of phase separation assay showing condensate formation of 5 µM full length MEG-3 incubated with poly-U RNA and/or MEX-5 as indicated. C. Violin plots showing condensate size and number for each experiment represented in B. The height of the plot shows the area of condensates in µm^2^. The width of the plot correlates to the proportion of condensates of that size. Numbers inside each violin plot are the total number of condensates pooled from three technical replicates for each condition.

### MEX-5 inhibits RNA-induced phase separation

To examine whether MEX-5 can affect MEG-3 phase separation, we purified the MEX-5 RNA-binding domain and C-terminus (aa236-468) as a His fusion (we were not able to obtain soluble full length MEX-5, Methods). We pre-incubated recombinant MEX-5 with poly-U in buffer for 30 minutes before adding MEG-3. We found that MEX-5 strongly inhibited MEG-3 phase separation induced by RNA (Fig 4B and C). Addition of excess RNA (3-fold increase) restored robust phase separation in the presence of MEX-5. These observations suggest that MEX-5 does not interfere with MEG-3 phase separation directly, but interferes with the ability of poly-U30 to induce phase separation.

### MEG-3__IDR__ forms a MEX-5-dependent gradient *in vivo* and can be stimulated to form granules by excess RNA

Our *in vitro* experiments indicate that MEG-3_IDR_ is sufficient to promote RNA binding and phase separation, but does so less efficiently than full length MEG-3 at low protein concentrations. To examine the behavior of MEG-3_IDR_ *in vivo*, we deleted the C-terminus of MEG-3 by CRISPR/Cas9 genome editing to generate a *meg-3* allele that only expresses MEG-3_IDR_ (Methods, Table S1). We found that, like full-length MEG-3, MEG-3_IDR_ is a cytoplasmic protein that redistributes into a posterior-rich gradient during polarization of the zygote (Fig 5A). Unlike MEG-3, however, MEG-3_IDR_ did not coalesce into prominent, micron-sized granules in zygotes (Fig 5A). Distinct MEG-3_IDR_ granules were observed starting in the 2-cell stage as MEG-3_IDR_ segregates into the progressively smaller P blastomeres (Fig 5A). In *mex-5/6(RNAi)* zygotes, MEG-3_IDR_ did not form a gradient and did not form granules (Fig 5B). These observations indicate that MEG-3_IDR_ is partially defective in granule formation, while remaining sensitive to MEX-5/6. The loss of *meg-3* activity was not a result of reduced expression as MEG-3_IDR_ was expressed at greater levels than wild-type MEG-3 (Fig S5).

**Figure 5.**
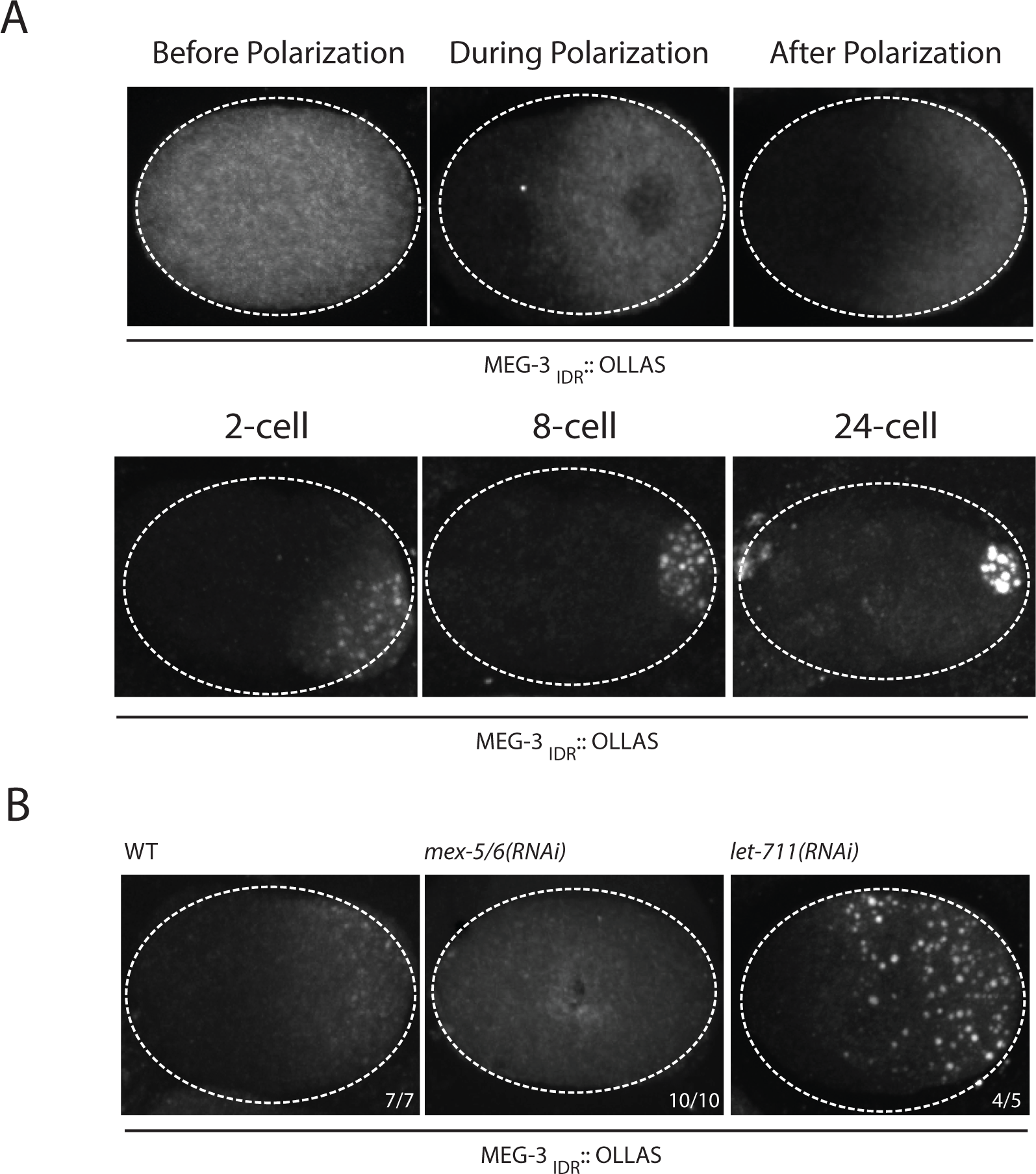
Coalescence of MEG-3_IDR_ can be stimulated by blocking mRNA turnover *in vivo*. A. Photomicrographs of fixed zygotes expressing MEG-3_IDR_ tagged with OLLAS epitope. Single cell (top) and late stage (bottom) zygotes are shown at indicated stages. B. Photomicrographs of fixed zygotes expressing MEG-3_IDR_ tagged with OLLAS epitope. Genotypes are indicated above each embryo (left to right: wild-type, *mex-5/6(RNAi)*, and *let-711* (*RNAi)*). Numbers indicate number of zygotes exhibiting phenotype shown / total number of zygotes examined. In 1/5 *let-711* (*RNAi)* zygotes, MEG-3_IDR_ formed granules but these were smaller and confined to the posterior half of the zygote, possibly due to incomplete depletion of *let-711*.

*In vitro*, the weaker phase separation properties of MEG-3_IDR_ at low protein concentrations can be stimulated by RNA. To determine whether excess RNA could also rescue granule formation by MEG-3_IDR_ in zygotes, we blocked maternal mRNA turnover by depleting LET-711 by RNAi. LET-711 is the scaffolding component of the CCF/NOT1 deadenylase, the main deadenylase that promotes mRNA turnover in oocytes and early embryos (DeBella, Hayashi et al. 2006, Nousch, Techritz et al. 2013). We found that MEG-3_IDR_ formed numerous micron-sized granules in *let-711(RNAi)* zygotes. The MEG-3_IDR_ granules and cytoplasmic gradient extended further towards the anterior compared to wild-type (Fig 5B). These observations suggest that, as we observed *in vitro*, excess RNA can overcome the inhibitory effects of MEX-5 and boost MEG-3 coalescence *in vivo*.

## DISCUSSION

P granule asymmetry in *C. elegans* zygotes is a text-book example of cytoplasmic partitioning (Strome and Wood 1983). In this study, we present evidence that P granule asymmetry is a direct consequence of an asymmetry in the distribution of the P granule scaffold MEG-3. MEG-3 localizes in a posterior-rich gradient under the control of the RNA-binding protein MEX-5, which localizes in a mirror-image, anterior-rich gradient. Our findings suggest that the MEG-3 gradient arises from an anterior-posterior gradient in RNA availability created by MEX-5. MEX-5 sequesters RNA in the anterior, which promotes MEG-3 phase separation in the posterior where MEX-5 concentration is low.

### MEG-3 is an RNA-binding protein that is stimulated by RNA to phase separate

MEG-3 contains a long N-terminal IDR but no recognizable RNA-binding domain. We have found that MEG-3 binds RNA (poly-U30) with nanomolar affinity *in vitro* (K_d,app_ = 32 nm). The MEG-3_IDR_ is essential for binding, but on its own binds with lower affinity (K_d,app_ = 460 nm). One possibility is that the MEG-3 IDR extends beyond the region predicted by IUPRED (Dosztanyi, Csizmok et al. 2005). The region immediately C-terminal scores close to the IUPRED cut-off (Wang, Smith et al. 2014), and may contribute to RNA binding. IDRs are over-represented among RNA-binding domains (Varadi, Zsolyomi et al. 2015, Castello, Fischer et al. 2016). Electrostatic interactions between positively-charged amino acids and the negatively-charged RNA backbone are often invoked as a possible mechanism for RNA binding by IDRs (Guo and Shorter 2015, Basu and Bahadur 2016). MEG-3 is rich in basic residues, but shows a strong preference for poly-U over poly-C and poly-G, suggesting that non-charged interactions are also involved.

Several recent studies have demonstrated that RNA-binding proteins containing IDRs phase separate in aqueous solutions (Guo and Shorter 2015). MEG-3 follows this paradigm: MEG-3 readily formed condensates within minutes of dilution from urea to an aqueous buffer (150mM NaCl). MEG-3 phase separation could be stimulated by RNA: addition of poly-U30 to the phase separation buffer increased the number of MEG-3 condensates especially at 1µM and lower protein concentrations. MEG-3_IDR_ behaved similarly to full-length MEG-3, except that MEG-3_IDR_ required higher concentrations of RNA to phase separate at low protein concentrations. Consistent with this *in vitro* behavior, MEG-3_IDR_ did not form large granules in wild-type zygotes, but could be induced to do so by blocking mRNA turnover. These observations suggest that the MEG-3_IDR_ confers on MEG-3 an intrinsic tendency for phase separation that is tunable by RNA. RNA-induced phase separation has also been observed for Whi3, a fungal RNA-binding protein, and for hnRNPA1, a stress granule protein (Lin, Protter et al. 2015, Zhang, Elbaum-Garfinkle et al. 2015). In these proteins, the IDR and RNA-binding domain are distinct and RNA-induced phase separation requires both domains. It will be interesting to determine whether the MEG-3_IDR_ in fact contains separable domains for phase separation and RNA binding.

### MEG-3 recruits PGL-1 assemblies: P granules contain multiple phases?

We showed previously that P granules are non-homogeneous organelles, and that MEG-3 overlaps but does not co-localize perfectly with PGL-3 in the granules (Wang, Smith et al. 2014). In this study, we show that MEG-3 assembles into posterior granules independently of PGL-1 and PGL-3, and is required (with MEG-4) to localize PGL-1 to posterior granules. MEG-3 binds PGL-1 *in vitro* and therefore could recruit PGL-1 to the granules directly (Wang, Smith et al. 2014). PGL-1 and PGL-3 are constitutive components of P granules that can assemble into cytoplasmic granules on their own and recruit other RNA-binding proteins (Hanazawa, Yonetani et al. 2011). Assembly of PGL granules was shown to require LAF-1, a DEAD-box RNA helicase that also localizes to P granules (Elbaum-Garfinkle, Kim et al. 2015). LAF-1 contains an IDR that is necessary and sufficient for phase separation *in vitro*. RNA does not stimulate LAF-1 phase separation, but reduces the viscosity of LAF-1 condensates (Elbaum-Garfinkle, Kim et al. 2015). Taken together, these observations raise the possibility that P granules comprise multiple phases each with distinct properties, specified by different assemblies of protein/RNA complexes that have affinity for one another but never fully mix. A similar model has been proposed for nucleoli, which exhibit a layered droplet organization specified by the biophysical properties of the components unique to each phase (Brangwynne, Eckmann et al. 2009). We suggest that MEG-3 and MEG-4 forms the main RNA-dependent phase of P granules and functions as a scaffold that recruits and/or stabilizes other phase-separated assemblies, including those assembled by LAF-1, PGL-1, and PGL-3.

### MEX-5 patterns MEG-3 by acting as an RNA sink

MEX-5 has been hypothesized to regulate P granule asymmetry by creating a supersaturation gradient of critical granule component(s) along the anterior-posterior axis of the zygote (Lee, Brangwynne et al. 2013). Our observations suggest that the critical component regulated by MEX-5 is RNA. MEX-5 binds RNA with nanomolar affinity (K_d,app_ = ∼29 nM, Pagano et al. 2007) and is 10-fold more abundant than MEG-3 in embryos (Fig S2A and S2B). In our *in vitro* phase separation assay, the MEX-5 RNA-binding domain was sufficient to inhibit RNA-induced phase separation of MEG-3. Similarly, *in vivo*, uniform MEX-5 was sufficient to inhibit MEG-3 granule assembly throughout the cytoplasm and this activity was disrupted by mutations that lower MEX-5’s affinity for RNA. Together, these results support a model where MEX-5 suppresses MEG-3 phase separation by removing RNA from the pool available to bind MEG-3. The MEX-5 gradient arises as a consequence of phosphorylation by PAR-1, which decreases the size of MEX-5 complexes and increases their diffusion rate (Griffin, Odde et al. 2011). One possibility is that phosphorylation by PAR-1 inhibits the formation of MEX-5/RNA complexes in the posterior cytoplasm leading to a local increase in “free” RNA available to bind MEG-3. A recent theoretical study proposed a similar “RNA sink” model, except that MEX-5 was hypothesized to regulate P granule asymmetry by suppressing the phase separation of another P granule protein, PGL-3 (Saha, Weber et al. 2016). PGL-3, however, is unlikely to be a critical MEX-5 target, since PGL-3 is not essential for granule assembly or asymmetry (Fig 1C and Fig S1B) and does not depend on its RNA-binding domain to localize to P granules (Hanazawa, Yonetani et al. 2011).

MEX-5’s high affinity for poly-U stretches, which are present in 91% of *C. elegans* 3’ UTRs (Pagano, Farley et al. 2007), suggests that MEX-5 interacts with most mRNAs in zygotes and thus could function as a general “RNA sink”. Consistent with this view, the MEX-5 gradient patterns the distribution of three other RNA-binding proteins that, like MEG-3, form posterior-rich gradients (Wu, Zhang et al. 2015). The observation that blocking mRNA turnover stimulates MEG-3 coalescence into macroscopic granules is consistent with the idea that the mRNA pool accessible to MEG-3 is limiting in zygotes. A limiting mRNA pool has also been suggested to regulate the balance of P bodies and stress granules in cells (Buchan and Parker 2009). We propose that regulated access to RNA, combined with RNA-induced phase separation of key scaffolding proteins, may be a general mechanism for controlling the formation of RNA granules in space and time.

## Materials and Methods

### CRISPR genome editing

*C. elegans* was cultured according to standard methods at 20^o^C (Brenner 1974). Genome editing was performed using CRISPR/Cas9 as described in Paix et al., 2015. Alleles used in this study are listed in Table S1.

### RNA mediated Interference (RNAi)

RNAi knock-down experiments were performed by feeding on HT115 bacteria (Timmons and Fire 1998). Feeding constructs were obtained from the Ahringer or Openbiosystems libraries and transformed into HT115 bacteria. pL4440 was used as a negative control empty feeding vector. Bacteria were grown at 37°C in LB + ampicillin (100 µg/mL) for 5 hours, induced with 5 mM IPTG for 45 minutes, plated on NNGM (nematode nutritional growth media) + ampicillin (100 µg/mL) + IPTG (1 mM), and grown overnight at room temperature. Embryos isolated by bleaching from gravid hermaphrodites were added to the RNAi plates and transferred to fresh plates as L4 larvae before examination of their progeny. All RNAi experiments were performed at 20^o^C.

### Protein Expression and Purification

All purifications were performed using an AKTA pure FPLC protein purification system (GE Healthcare).

Purification of MEG-3 and MEG-3_IDR_: MEG-3 (aa1-862), MEG-3_IDR_ (aa1-544) fused to an N-terminal 6XHis tag in pET28a were expressed in Rosetta (DE3) cells at 16°C in LB + ampicillin (100 µg/mL) to an OD600 of ∼0.4 and induced with 0.4 mM isopropyl β-D-1-thiogalactopyranoside at 16° C for 16 hours. Cells were resuspended in Buffer A (20 mM HEPES, 500 mM NaCl, 20 mM Imidazole, 10% (vol/vol) glycerol, 1% Triton-X100, 6M Urea, pH7.4) with added protease inhibitors and TCEP, lysed by sonication, spun at 13,000 rpm for 25 minutes, and incubated overnight at 4C. Lysate was passed over a His Prep FF 16/10 column (GE Healthcare). Bound protein was washed with Buffer B (20 mM HEPES, pH 7.4, 1M NaCl, 25 mM Imidazole, 10% (vol/vol) glycerol, 6M urea, 6 mM βME) and eluted in Buffer C (20 mM HEPES, pH 7.4, 1M NaCl, 250 mM Imidazole, 10% (vol/vol) Glycerol, 6M Urea, 6 mM βME). After each purification, aliquots of the peak elution fraction were run on 4-12% Bis Tris gels, and stained with Simply Blue Safe Stain (ThermoFisher). Proteins were concentrated to a final concentration of 100-320 µM in elution Buffer C. For use in RNA binding assays, proteins were dialyzed into storage buffer B (25 mM Hepes, pH 7.4, 1M NaCl, 10% (vol/vol) glycerol) and stored at -80° C.

Purification of MEG-3_Cterm_: MEG-3(545-862) was purified as above and also natively using the same protocol without urea. MEG-3_Cterm_ purified under native conditions was soluble in aqueous buffer even at high concentrations (>1uM) and was used for RNA-binding assays.

Purification of MEX-5: MEX-5(aa236-468) was purified as an N-terminal 6xHis:MBP fusion expressed in Rosetta (DE3) competent cells. Cells were grown at 37° C in LB + ampicillin (100µg/mL) to an OD600 of ∼0.4, before induction with 0.2 mM isopropyl β-D-1-thiogalactopyranoside and 100 µM zinc acetate at 16° C for 16 hours. Cells were resuspended in lysis buffer (20 mM Tris-HCl, pH 8.3, 200 mM NaCl, 20 mM imidazole, 10% (vol/vol) glycerol, 1 mM TCEP, 100 µM zinc acetate, Roche complete EDTA-free protease inhibitor tablet), lysed by sonication and pelleted at 10,000 rcf for 15 minutes. The supernatant was passed over a HisTrap HP column (GE Healthcare) and washed with wash buffer (20 mM Tris-HCl, pH8.3, 800 mM NaCl, 20 mM imidazole, 10% (vol/vol) glycerol, 100 µM zinc acetate,1 mM TCEP). Column was eluted using elution buffer (20 mM HEPES, pH 8.3, 500 mM NaCl, 250 mM imidazole, 10% (vol/vol) glycerol, 100 µM zinc acetate). Elution fractions were pooled and run over a HiTrap Heparin HP column (GE Healthcare). Column was then washed in wash buffer B (20 mM Hepes, pH 8.4, 200 mM NaCl) and eluted using a gradient of wash buffer B and elution buffer B (20 mM Hepes, pH 8.4, 1.5 M NaCl, 100 µM zinc acetate). Elutions were pooled and dialyzed into storage buffer as in Pagano et al., 2007 (20 mM Tris, pH 8.3, 20 mM NaCl, 100 µM zinc acetate, 10% (vol/vol) glycerol). Protein concentration was determined by measuring absorbance at 280 nm as in Pagano et al., 2007 and stored at -80°C.

### Immunostaining

Adult worms were placed into M9 salt solution on epoxy autoclavable slides (thermo-fisher) and squashed with a coverslip to extrude embryos. Slides were frozen by laying on pre-chilled aluminum blocks for 20 minutes (chilled using dry ice). Embryos were permeabilized by freeze-cracking (removal of coverslips from slides) followed by incubation in methanol at -20°C for >15 minutes, and in acetone (pre-chilled at -20^o^C) at room temperature for 10 minutes. Slides were blocked in PBS-Tween (0.1%) BSA (0.5%) for 15 minutes x 2, and incubated with 50 ul primary antibody overnight at 4°C in a humid chamber. Antibody dilutions (in PBST/BSA): K76 (1:10, DSHB), Rat α OLLAS-L2 (1:200, Novus Biological), mouse α FLAG (1:500, Sigma). Secondary antibodies were applied for 2 hours at room temperature.

### Confocal Microscopy

Fluorescence microscopy was performed using a Zeiss Axio Imager with a Yokogawa spinning-disc confocal scanner. Images were taken and stored using Slidebook v 6.0 software (Intelligent Imaging Innovations) using a 63x objective. For live imaging, embryos were dissected from adult hermaphrodites in M9 salt solution and mounted onto 3% agarose pads. All embryo images are z stack maximum projections using a z step size of 1 µm, spanning the entire width of the embryo.

### Quantification of MEG-3::meGFP fluorescence from confocal images

Equally normalized time-lapse images were quantified using Slidebook v 6.0. Average fluorescence intensity relative to area of anterior (60%) and posterior (40%) of zygote were quantified and average fluorescence intensity relative to area of background (outside of zygote) was subtracted from each of these values. For each time-point, anterior and posterior fluorescence were expressed as fractions of total fluorescence and then normalized to T_0_ (14 minutes prior to mitosis). Final values represent average of three embryos. Error bars show standard deviation of the mean.

### Electrophoretic Mobility Shift Assay (EMSA)

EMSA were carried out as described in Pagano et al. 2007. Reactions consisted of 50 nM 3’ Fluorescein-labeled RNA oligonucleotides (Dharmacon -GE Lifesciences) incubated with protein for 2 hours or more at room temperature or for 30 minute followed by 2 hour incubation with unlabeled competitor. Samples were run on 1% agarose gel in 1x TAE. Gels were scanned immediately using typhoon FLA-9500 with blue laser at 473 nm.

### Fluorescence Polarization Assay

Equilibration reactions were performed using same protocol as for EMSAs. Reactions were transferred to 384 well microplates (Greiner Bio-One). The apparent fluorescence polarization was determined using a Clariostar monochromator microplate reader with fluorescein-sensitive filters and polarizers. Polarization values were normalized relative to saturation polarization value. For each experiment, values of three reads were averaged. Average values and standard errors from at least three technical replicates were calculated and plotted against each protein concentration. These data were fit to a quadratic equation (equation 1, where b is the base polarization, m is the maximum polarization, R is the labeled nucleic acid concentration, and P is the total protein concentration) as in (Pagano, Clingman et al. 2011), to calculate the apparent dissociation constant. The reported values are the dissociation constants calculated using the polarization values averaged from all technical replicates. The reported errors are standard error values calculated from the dissociation constant of each individual technical replicate.

Equation 1

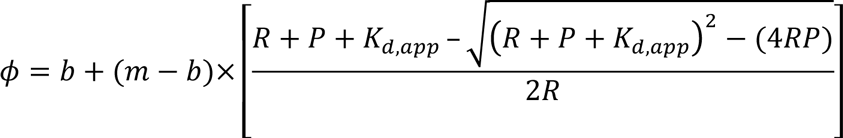

### Phase Separation Assay

His::tagged MEG-3 fusions were quickly diluted out of urea into condensation Buffer (25mM HEPES, pH7.4, NaCl adjusted to a final concentration of 150mM) in the presence and absence of poly-U30 RNA. Dilutions were performed by adding buffer to protein in low-binding siliconized Eppendorf tubes and mixing briefly by pipetting. The reaction was either spun at 3000 rpm for 2 minutes or transferred directly into 35mm glass bottom dish (Cat. No. P35G-1.5-14-C MatTek Corp) for imaging. Phase separation assays with MEX-5 were performed by pre-incubating 3.5µM 6xHis::MBP::MEX-5 protein (dialyzed into condensation buffer) with condensation buffer and poly-U30 RNA for 30 minutes before diluting in MEG-3. Differential interference contrast (DIC) images were obtained on an Olympus inverted microscope, using a 100X objective. Images were taken and stored using Slidebook v 4.0 and 5.0. All images are a single focal plane focused on the slide surface. For each phase separation experiment, we took three separate images of an 80 x 80 micron field and counted the condensates using Image J64. To recognize condensates, background was subtracted from each image using a rolling ball radius of 10 pixels, a pixel brightness threshold was set to 15-255. Remaining pixels were smoothed 3 times and size and number of objects greater than 0.032 µm^2^ were quantified. Quantifications were manually verified for each image used. At least 3 technical replicates were quantified for each condition. Average number of condensates and S.E.M were calculated using all technical replicates.

### Western Blots

Western blots were performed by running worm lysates on 7% Tris Acetate SDS PAGE precast gels (Bio-Rad). Protein was transferred to nitrocellulose membrane which was pre-blocked in 5% Milk diluted in PBS-Tween (0.1%) for 5 minutes (3 times). Membrane was then incubated with Primary antibody for at least 18 hours at 4° C or 2 hours at room temperature. Membranes were washed and blocked in 5% milk for 5 minutes (3 times) and incubated with secondary HRP conjugated antibody for 45 minutes at room temperature. Membranes were washed in 5% milk for 5 minutes (2 times) and PBST for 5 minutes (1 time). Membranes were then exposed to ECL substrate for 1 minute and then exposed to film. Primary antibody dilutions (in 5% Milk PBST): Rat α OLLAS-L2 (1:1000, Novus Biological), Mouse α Tubulin (1:1000, Sigma).

## Acknowledgements

We thank the CGC (USA) for strains, the Berger lab (JHU) for reagents and use of their microplate reader, Alex Paix for the *glh(ax3064)* allele, and the JHMI microscope facility for instruction. Research in the Seydoux lab is supported by R01 HD37047 from the National Institute of Health. G. Seydoux is an Investigator of the Howard Hughes Medical Institute.

## Supplemental Figures

**Figure S1.**
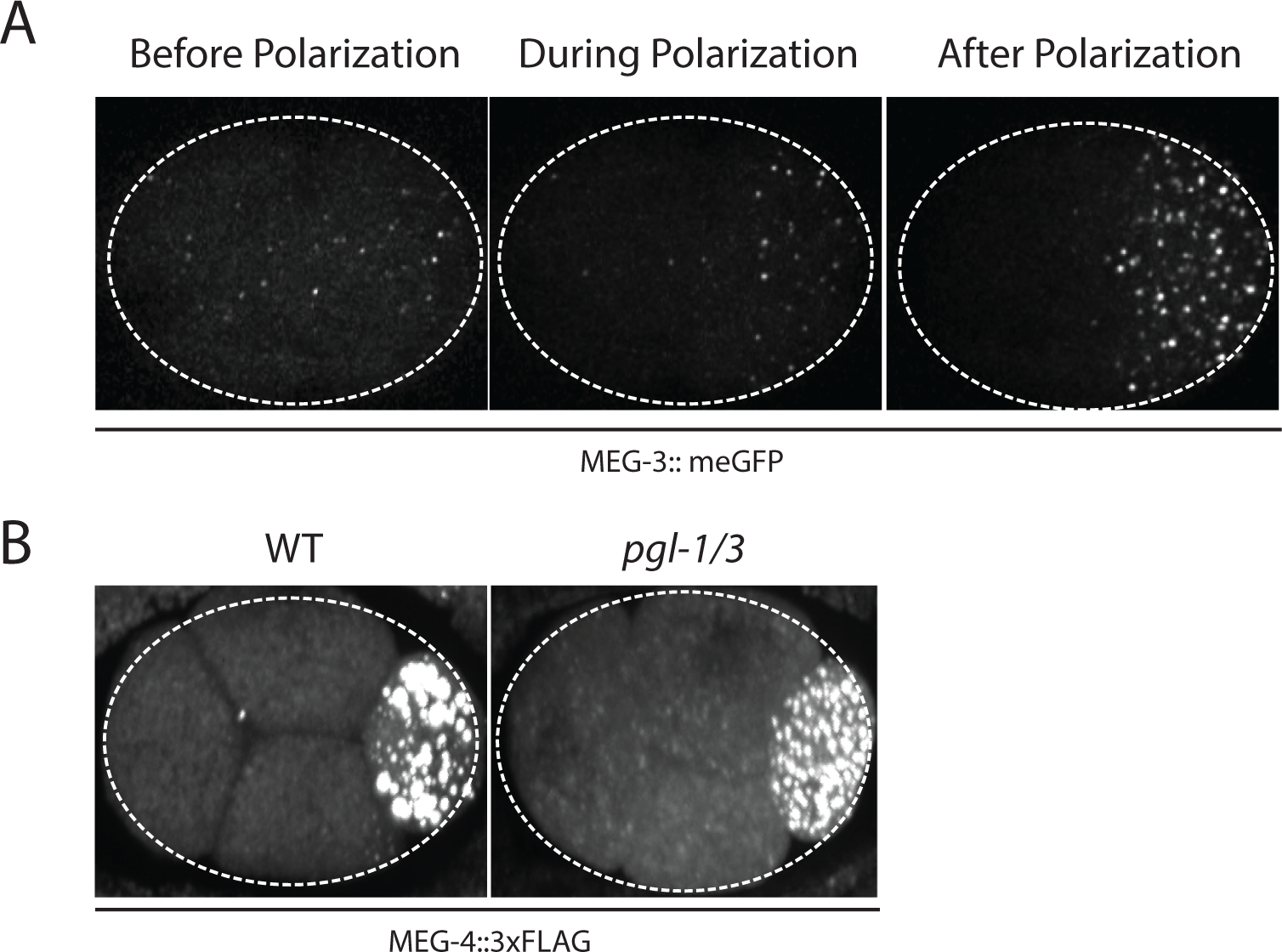
MEG-3 localizes before PGL-1 and does not require PGL-1 and PGL-3 to assemble granules. A. Photomicrographs of live MEG-3::meGFP in zygotes at three different stages: before polarization (pronuclear formation), during polarization (pronuclear migration) and after polarization (mitosis). These and similar images taken from 3 embryos at 14 time points were used to quantify MEG-3::meGFP fluorescence levels over time as shown in Fig 1B. B. Photomicrographs of fixed *pgl-1/3* zygotes stained with anti-FLAG to show MEG-4 localization at the 4-cell stage. *pgl-1/3* zygotes were derived from *pgl-3(bn104); meg-4(ax2080^FLAG tag^)* hermaphrodites treated with *pgl-1* RNAi.

**Figure S2.**
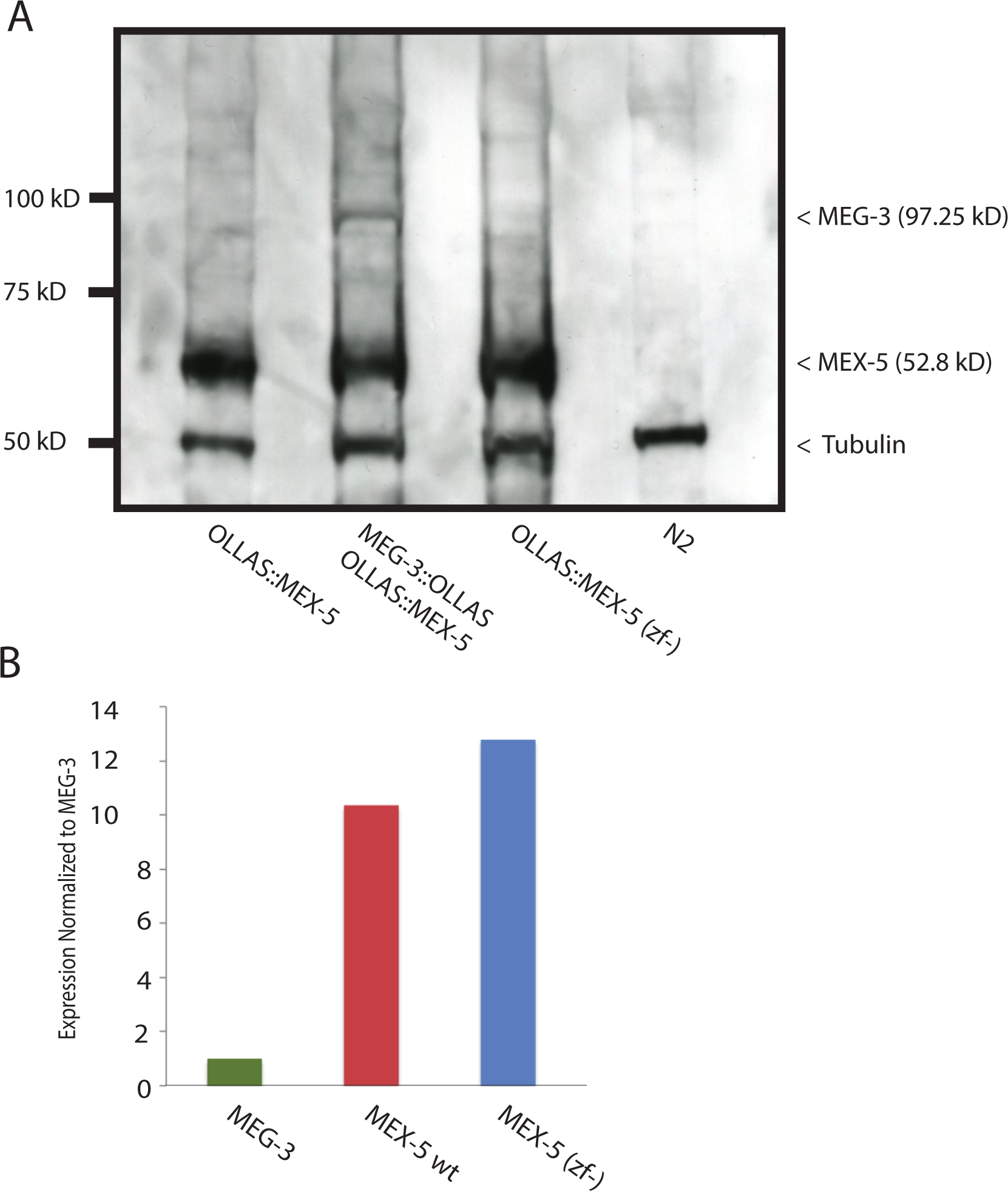
Expression of MEX-5, and MEX-5 (ZF-), MEG-3, and MEG-3_IDR_. A. Western blot of embryo lysates co-blotted with α OLLAS and α tubulin (loading control). Expected sizes for protein-epitope fusions are indicated on the right by a < symbol. Lysates are shown for (left to right) MEX::OLLAS, MEX-5::OLLAS; MEG-3::OLLAS, and MEX-5::OLLAS (ZF-) and control lysates with no epitope expression (N2). B. Bar graph showing relative expression of MEG-3 and MEX-5. Western blot in figure S2A was quantified using Image J. MEX-5 is 10X more abundant than MEG-3.

**Figure S3.**
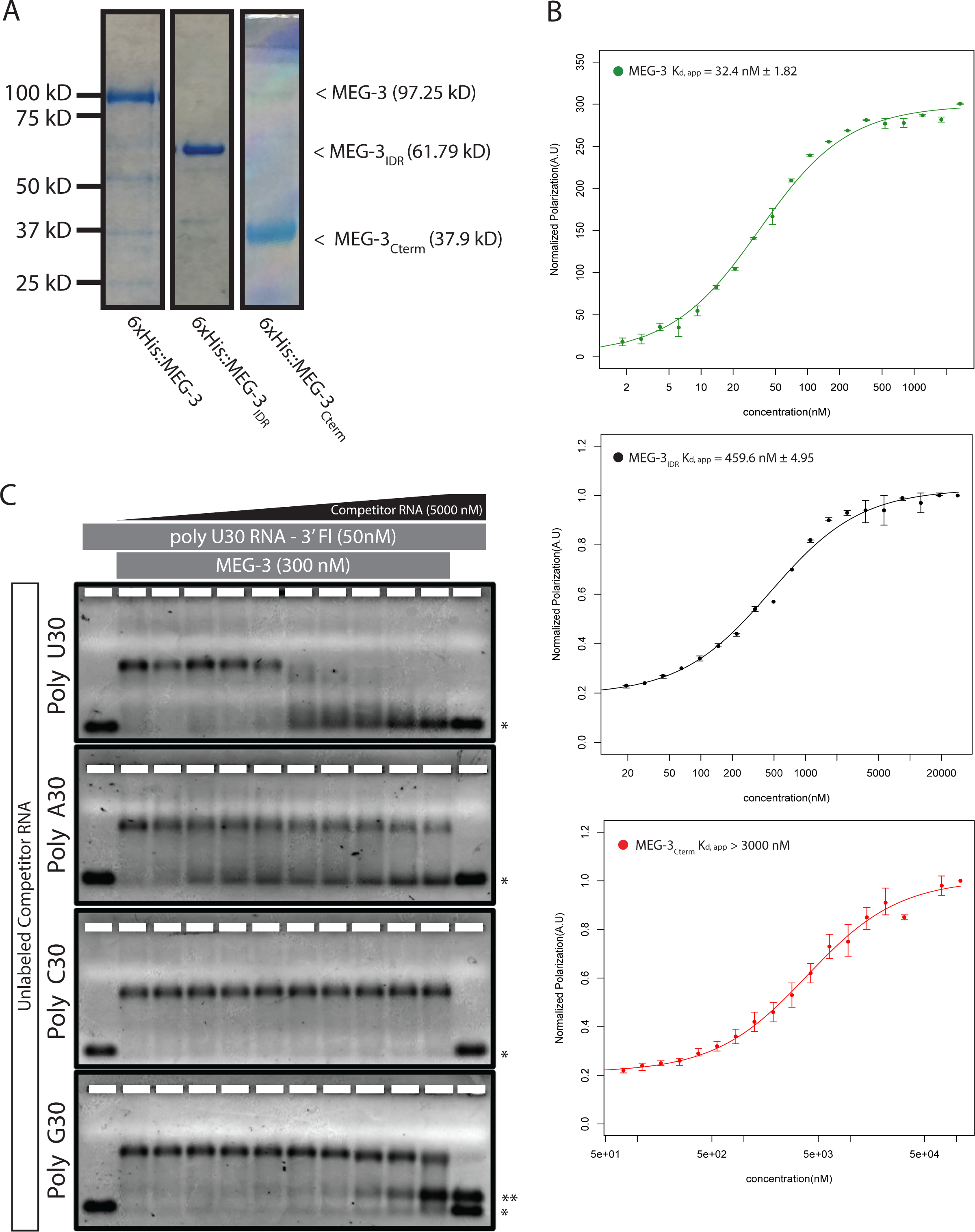
MEG-3 RNA binding. A. Coomassie stained gel showing His-tagged MEG-3 proteins. Expected sizes are indicated on the right. Each lane is from a separate gel. B. RNA-RNA competition EMSA. 300nM MEG-3 (300 nM) was incubated with 50nM poly-U30 and increasing amounts of unlabeled 30-mer RNAs as indicated the left of each panel. The first lane of each gel contains only labeled poly-U RNA. The last lane of each gel contains poly-U RNA and the highest concentration of unlabeled competitor RNA (5000 nM). The unbound poly-U RNA is denoted by an asterisk (*). Double asterisks (**) in the poly-G assay denote a band that arises due to an interaction between poly-G and poly-U (independent of MEG-3). C. Fluorescence Polarization of polyuridine by MEG-3 (expanded from Fig. 3). Fluorescence polarization values normalized relative to saturation are shown for full length MEG-3 (top, green), MEG-3_IDR_ (middle, black), and MEG-3_Cterm_(bottom, red). A fit of the polarization as function of protein concentration is plotted and used to calculate the given K_d,app_. Error bars report S.E.M.

**Figure S4.**
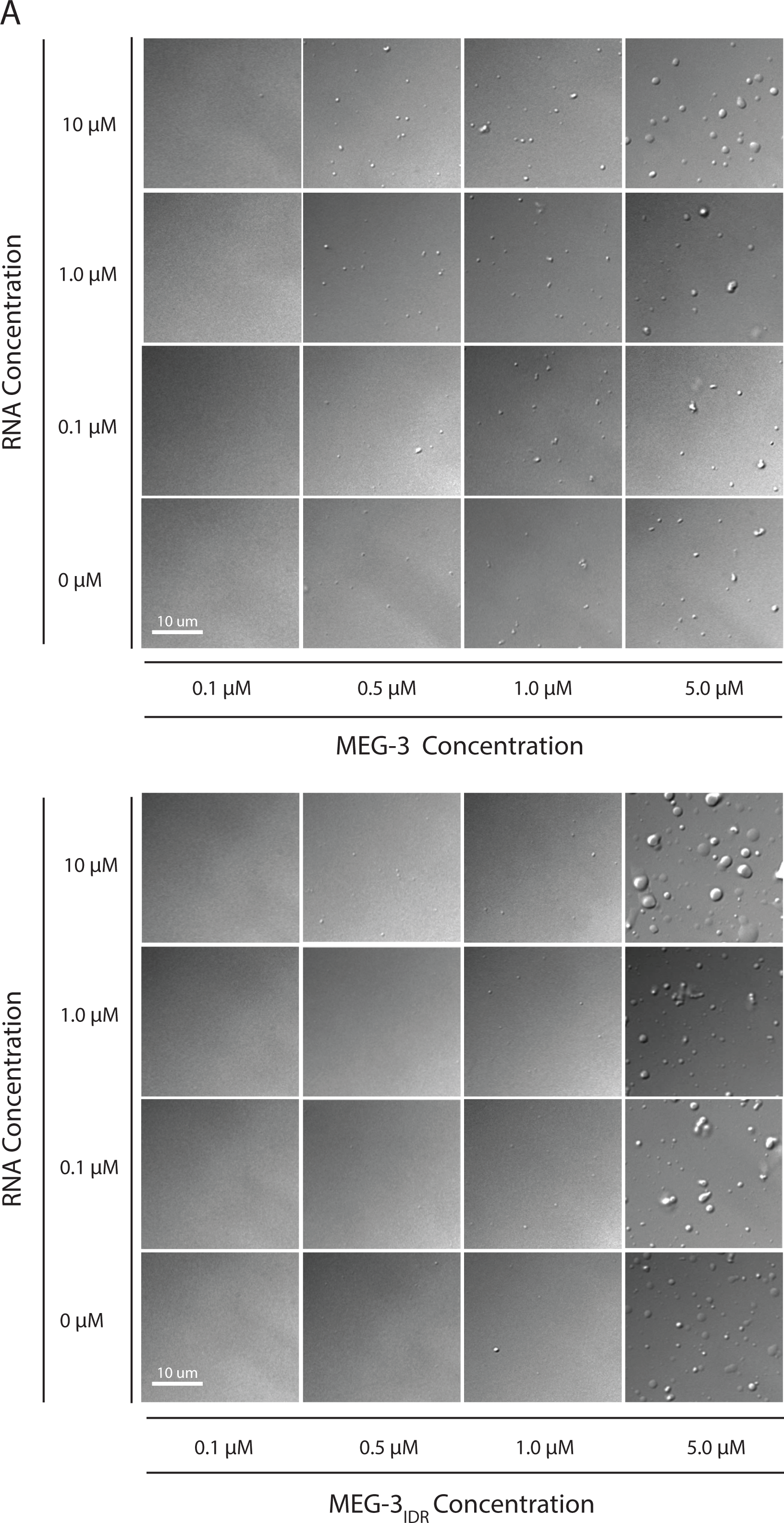
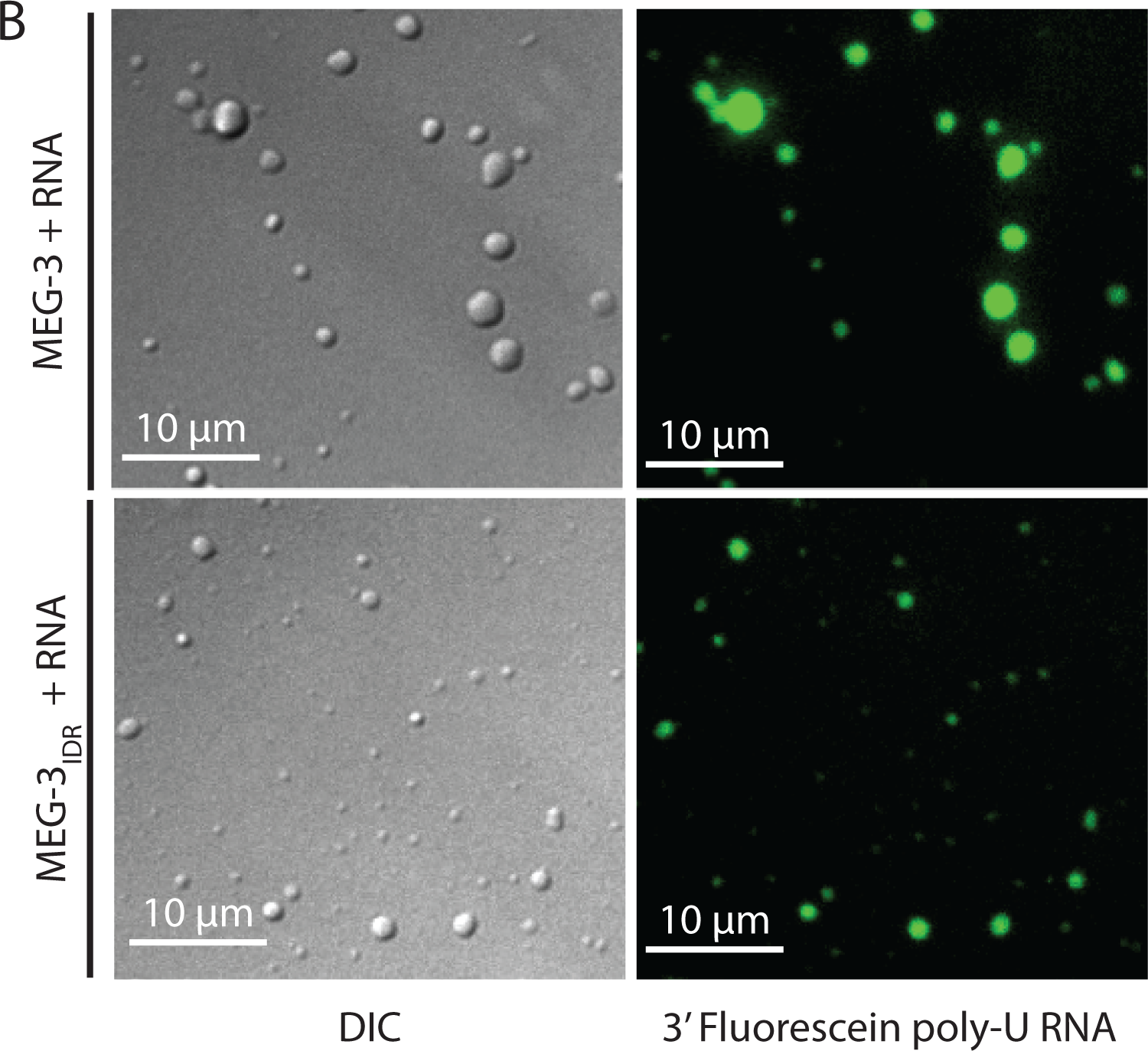
Phase Separation assay. A. DIC images of MEG-3 solutions in the presence of increasing concentrations of protein and RNA. MEG-3 was diluted into phase separation buffer (25mM HEPES, pH7.4, NaCl adjusted to a final concentration of 150mM) at room temperature. The solution was transferred to a glass bottom dish and the dish surface was photographed using an inverted DIC microscope 10 minutes after the initial dilution (room temperature). B. Same as above except 3’ fluorescein labeled poly-U RNA was included in the buffer. DIC images (left) show the MEG-3 condensates and fluorescence images (488 channel) show the 3’ fluorescein labeled poly-U RNA concentrated in the condensates.

**Figure S5.**
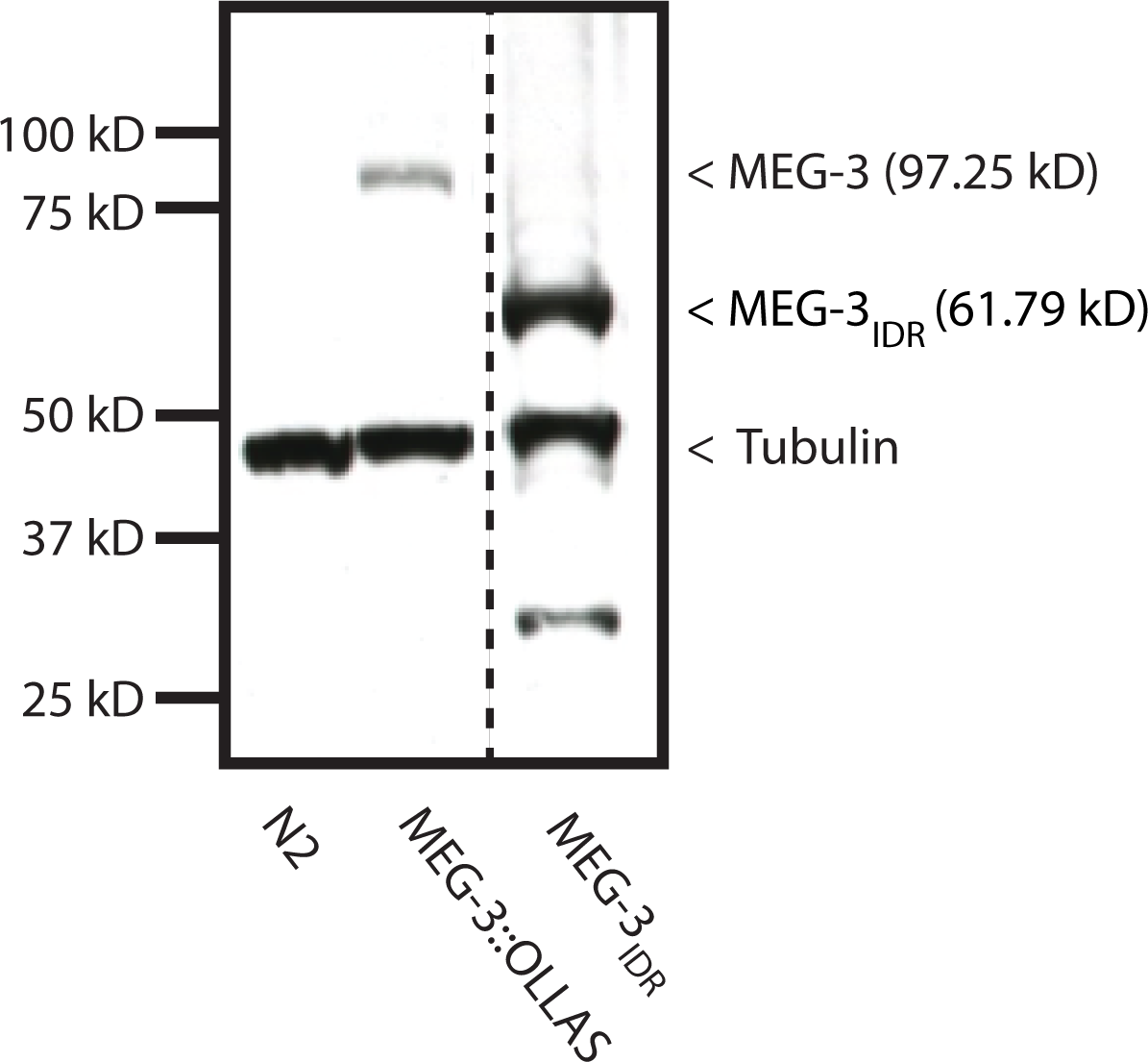
MEG-3_IDR_*in vivo*. Western blot of mixed-stage embryo lysates co-blotted with α OLLAS and α tubulin. Expected sizes for protein-epitope fusions are indicated on the right by a < symbol. Dashed line indicates a break in the original gel. Genotype is indicated on the bottom. The MEG-3 _IDR_ is present at higher level than MEG-3, likely due to the fact that unlike MEG-3, MEG-3 _IDR_ is not turned over in somatic cells (>Fig. 5). The significance of this difference is not known.

**Table S1.**
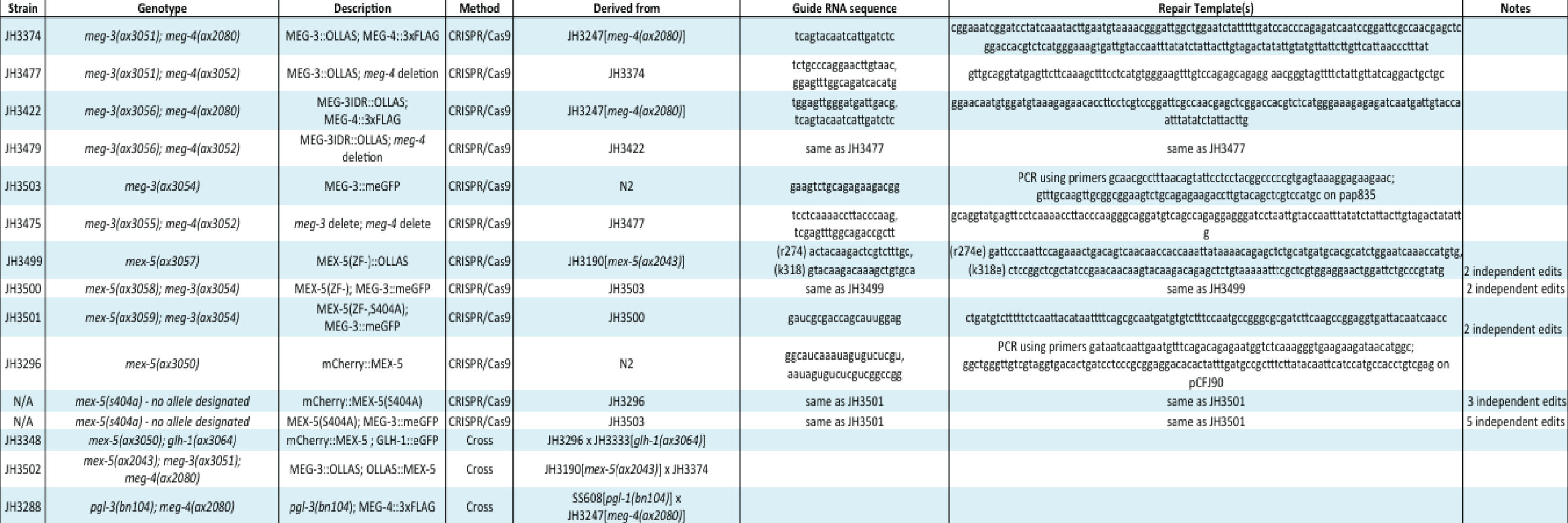
Strains used in this study. All strains were generated in this study by genome editing or crossing. No transgenic lines were used. Independent edits displayed the same phenotypes. The *mex-5(S404A)* lines could not be maintained due to semi-dominant maternal-effect sterility (91.6%) and recessive maternal-effect lethality (100%).

## References

Anderson, P. and N. Kedersha (2006). “RNA granules.” J Cell Biol 172(6): 803–808.

Basu, S. and R. P. Bahadur (2016). “A structural perspective of RNA recognition by intrinsically disordered proteins.” Cell Mol Life Sci.

Brangwynne, C.P., et al. (2009). “Germline P granules are liquid droplets that localize by controlled dissolution/condensation.” Science 324(5935): 1729–1732.

Brenner, S. (1974). “The genetics of Caenorhabditis elegans.” Genetics 77(1): 71–94.

Buchan, J. R. and R. Parker (2009). “Eukaryotic stress granules: the ins and outs of translation.” Mol Cell 36(6): 932–941.

Castello, A., et al. (2016). “Comprehensive Identification of RNA-Binding Domains in Human Cells.” Mol Cell 63(4): 696–710.

Courchaine, E.M., et al. (2016). “Droplet organelles?” EMBO J 35(15): 1603–1612.

DeBella, L.R., et al. (2006). “LET-711, the Caenorhabditis elegans NOT1 ortholog, is required for spindle positioning and regulation of microtubule length in embryos.” Mol Biol Cell 17(11): 4911–4924.

Dosztanyi, Z., et al. (2005). “IUPred: web server for the prediction of intrinsically unstructured regions of proteins based on estimated energy content.” Bioinformatics 21(16): 3433–3434.

Elbaum-Garfinkle, S., et al. (2015). “The disordered P granule protein LAF-1 drives phase separation into droplets with tunable viscosity and dynamics.” Proc Natl Acad Sci U S A 112(23): 7189–7194.

Gallo, C.M., et al. (2010). “Cytoplasmic partitioning of P granule components is not required to specify the germline in C. elegans.” Science 330(6011): 1685–1689.

Griffin, E.E., et al. (2011). “Regulation of the MEX-5 gradient by a spatially segregated kinase/phosphatase cycle.” Cell 146(6): 955–968.

Guo, L. and J. Shorter (2015). “It’s Raining Liquids: RNA Tunes Viscoelasticity and Dynamics of Membraneless Organelles.” Mol Cell 60(2): 189–192.

Hanazawa, M., et al. (2011). “PGL proteins self associate and bind RNPs to mediate germ granule assembly in C. elegans.” J Cell Biol 192(6): 929–937.

Kato, M., et al. (2012). “Cell-free formation of RNA granules: low complexity sequence domains form dynamic fibers within hydrogels.” Cell 149(4): 753–767.

Lee, C.F., et al. (2013). “Spatial organization of the cell cytoplasm by position-dependent phase separation.” Phys Rev Lett 111(8): 088101.

Li, P., et al. (2012). “Phase transitions in the assembly of multivalent signalling proteins.” Nature 483(7389): 336–340.

Lin, Y., et al. (2015). “Formation and Maturation of Phase-Separated Liquid Droplets by RNA-Binding Proteins.” Mol Cell 60(2): 208–219.

Motegi, F. and G. Seydoux (2013). “The PAR network: redundancy and robustness in a symmetry-breaking system.” Philos Trans R Soc Lond B Biol Sci 368(1629): 20130010.

Nott, T.J., et al. (2015). “Phase transition of a disordered nuage protein generates environmentally responsive membraneless organelles.” Mol Cell 57(5): 936–947.

Nousch, M., et al. (2013). “The Ccr4-Not deadenylase complex constitutes the main poly(A) removal activity in C. elegans.” J Cell Sci 126(Pt 18): 4274–4285.

Pagano, J.M., et al. (2011). “Quantitative approaches to monitor protein-nucleic acid interactions using fluorescent probes.” RNA 17(1): 14–20.

Pagano, J.M., et al. (2007). “Molecular basis of RNA recognition by the embryonic polarity determinant MEX-5.” J Biol Chem 282(12): 8883–8894.

Paix, A., et al. (2015). “High Efficiency, Homology-Directed Genome Editing in Caenorhabditis elegans Using CRISPR-Cas9 Ribonucleoprotein Complexes.” Genetics 201(1): 47–54.

Pitt, J.N., et al. (2000). “P granules in the germ cells of Caenorhabditis elegans adults are associated with clusters of nuclear pores and contain RNA.” Dev Biol 219(2): 315–333.

Saha, S., et al. (2016). “Polar Positioning of Phase-Separated Liquid Compartments in Cells Regulated by an mRNA Competition Mechanism.” Cell.

Schubert, C. M., et al. (2000). “MEX-5 and MEX-6 function to establish soma/germline asymmetry in early C.elegans embryos.” Mol Cell 5(4): 671–682.

Strome, S. and W. B. Wood (1983). “Generation of asymmetry and segregation of germ-line granules in early C.elegans embryos.” Cell 35(1): 15–25.

Timmons, L. and A. Fire (1998). “Specific interference by ingested dsRNA.” Nature 395(6705): 854.

Updike, D. and S. Strome (2010). “P granule assembly and function in Caen

Varadi, M., et al. (2015). “Functional Advantages of Conserved Intrinsic Disorder in RNA-Binding Proteins.” PLoS One 10(10): e0139731.

Voronina, E., et al. (2011). “RNA granules in germ cells.” Cold Spring Harb Perspect Biol 3(12).

Wang, J.T., et al. (2014). “Regulation of RNA granule dynamics by phosphorylation of serine-rich, intrinsically disordered proteins in C. elegans.” Elife 3:e04591.

Weber, S. C. and C. P. Brangwynne (2012). “Getting RNA and protein in phase.” Cell 149(6): 1188–1191.

Wu, Y., et al. (2015). “Coupling between cytoplasmic concentration gradients through local control of protein mobility in the Caenorhabditis elegans zygote.” Mol Biol Cell 26(17): 2963–2970.

Zhang, H., et al. (2015). “RNA Controls PolyQ Protein Phase Transitions.” Mol Cell 60(2): 220–230.

